# DeepHE: Accurately Predicting Human Essential Genes based on Deep Learning

**DOI:** 10.1101/2020.02.14.950048

**Authors:** Xue Zhang, Wangxin Xiao, Weijia Xiao

## Abstract

**Motivation:** Accurately predicting essential genes using computational methods can greatly reduce the effort in finding them via wet experiments at both time and resource scales, and further accelerate the process of drug discovery. Several computational methods have been proposed for predicting essential genes in model organisms by integrating multiple biological data sources either via centrality measures or machine learning based methods. However, the methods aiming to predict human essential genes are still limited and the performance still need improve. In addition, most of the machine learning based essential gene prediction methods are lack of skills to handle the imbalanced learning issue inherent in the essential gene prediction problem, which might be one factor affecting their performance.

**Results:** We proposed a deep learning based method, DeepHE, to predict human essential genes by integrating features derived from sequence data and protein-protein interaction (PPI) network. A deep learning based network embedding method was utilized to automatically learn features from PPI network. In addition, 89 sequence features were derived from DNA sequence and protein sequence for each gene. These two types of features were integrated to train a multilayer neural network. A cost-sensitive technique was used to address the imbalanced learning problem when training the deep neural network. The experimental results for predicting human essential genes showed that our proposed method, DeepHE, can accurately predict human gene essentiality with an average AUC higher than 94%, the area under precision-recall curve (AP) higher than 90%, and the accuracy higher than 90%. We also compared DeepHE with several widely used traditional machine learning models (SVM, Naïve Bayes, Random Forest, Adaboost). The experimental results showed that DeepHE greatly outperformed the compared machine learning models.

**Conclusions:** We demonstrated that human essential genes can be accurately predicted by designing effective machine learning algorithm and integrating representative features captured from available biological data. The proposed deep learning framework is effective for such task.

**Availability and Implementation:** The python code will be freely available upon the acceptance of this manuscript at https://github.com/xzhang2016/DeepHE.

## 1 Introduction

Essential genes are a subset of genes which are indispensable to the survival or reproduction of a living organism. The prediction of gene essentiality is very important for understanding the minimal requirements of an organism, identifying disease genes, and finding new drug targets. The discovery of essential genes via wet-lab experimental methods are often time-consuming, laborious, and costly. With the accumulation of gene essentiality data in some model organisms and human cell lines, many computational methods have been proposed to predict essential genes by exploring the correlations between gene essentiality and all sorts of biological information.

One focus in this direction is network based centrality measures. Many studies have demonstrated that highly connected proteins in a protein-protein interaction (PPI) network are more likely to be essential than those of low connections. Although the so-called centrality-lethality rule has been observed in several species, the prediction accuracy is very low for predicting gene essentiality solely based on each of these network topological features. One reason is that the existing PPI networks are not complete and very noisy. The other reason might be the fact that gene essentiality is expected to be affected by multiple biological factors which cannot be fully captured by network topological features. To improve the prediction accuracy, several new centrality measures have been proposed by combining topological properties with other biological information. For example, CoEWC integrated network topological property with gene expression data to capture the common features of essential proteins in both date hubs and party hubs, and showed significant performance improvement compared to methods only based on PPI networks [1]. Zhang et al. proposed an ensemble framework based on gene expression data and PPI networks, which can significantly improve the prediction accuracy of common used centrality measures [2]. Zhang et al. also proposed an integrated method, OGN, by combining network topological properties, the probability of co-expression with the neighboring proteins, and the orthologs in reference organisms [3]. Li et al. proposed GOS [4] by integrating gene expression, orthology, subcellular localization and PPI networks to predict gene essentiality. UDoNC combined the domain features with the topological properties of PPI networks to predict protein essentiality [5]. Centrality measure based methods predict gene essentiality by a scalar score derived whether from biological network or by integrating multiple data sources, which have limited power for accurately identifying all essential genes. More details about centrality measures for predicting essential genes/proteins can be found in a recent review [6].

The other focus is using machine learning to integrate multiple features derived from different biological data sources to predict gene essentiality. Zhang et al. provided a comprehensive review for gene essentiality predicting methods based on machine learning and network topological features, and pointed out the challenges and potential research directions [7]. As shown in [7], most of the proposed machine learning based predicting methods were evaluated in model organisms. In addition, the traditional machine learning methods were used to predict gene essentiality. Recently, Guo et al. used SVM (Support Vector Machines) to predict human gene essentiality based on the *λ*-interval Z curve derived features from nucleotide sequence data [8]. Zeng et al. used deep learning method to predict gene essentiality by integrating gene expression data, subcellular localization data, and PPI networks together, and tested it on *S. cerevisiae* [9]. Hasan et al. used a six hidden-layers neural network to predict gene essentiality in microbes based on sequence data [10].

Recently, human essential genes were identified in several human cancer cell lines using CRISPR-Cas9 and gene-trap technology [11-13]. These identified essential genes provided a clear definition of the requirements for sustaining the basic cell activities of individual human tumor cell types, and can be regarded as targets for cancer treatment [14]. These essential gene datasets together with other available biological data sources enable us to test one important and interesting assumption that human gene essentiality might be accurately predicted using computational methods. Although many previous studies showed that features derived from experimental omics data are useful to predict gene essentiality, such experimental omics data are often unavailable for under studied organisms. In this paper, we proposed a deep learning based method to predict human gene essentiality by using features derived from sequence data, which is therefore easily ready to be used for predicting essential genes in other organisms. In addition, in order to improve the performance of the proposed method, we also explored features automatically learned by using a deep learning embedding method from human protein interaction network. We showed that each of the two types of features can train a classifier with acceptable prediction performance, and the integration of these features further improves the prediction accuracy.

## 2 The Proposed Deep Learning Framework

Figure 1 gives the overall architecture of the proposed deep learning framework, DeepHE. It mainly consists of two parts, feature extraction and classification. It takes two types of data as input, the sequence data and PPI network. At the feature extraction level, several sequence features for each gene were extracted from the nucleotide sequence and protein sequence data. In addition, an embedding method, node2vec [15], was used to learn the semantic features for each gene from the PPI network. The classification module consists of several fully connected hidden layers and an output layer. All hidden layers utilized an excellent activation function, ReLU (Rectified Linear Unit), as their activation functions, and used dropout parameter to prevent overfitting. After the hidden layers, a fully connected output layer used softmax as its activation function. Considering the skewed distribution nature of human essential gene prediction problem, we explored a cost-sensitive technique to address the imbalanced learning issue when training the classifier.

**Figure 1.**
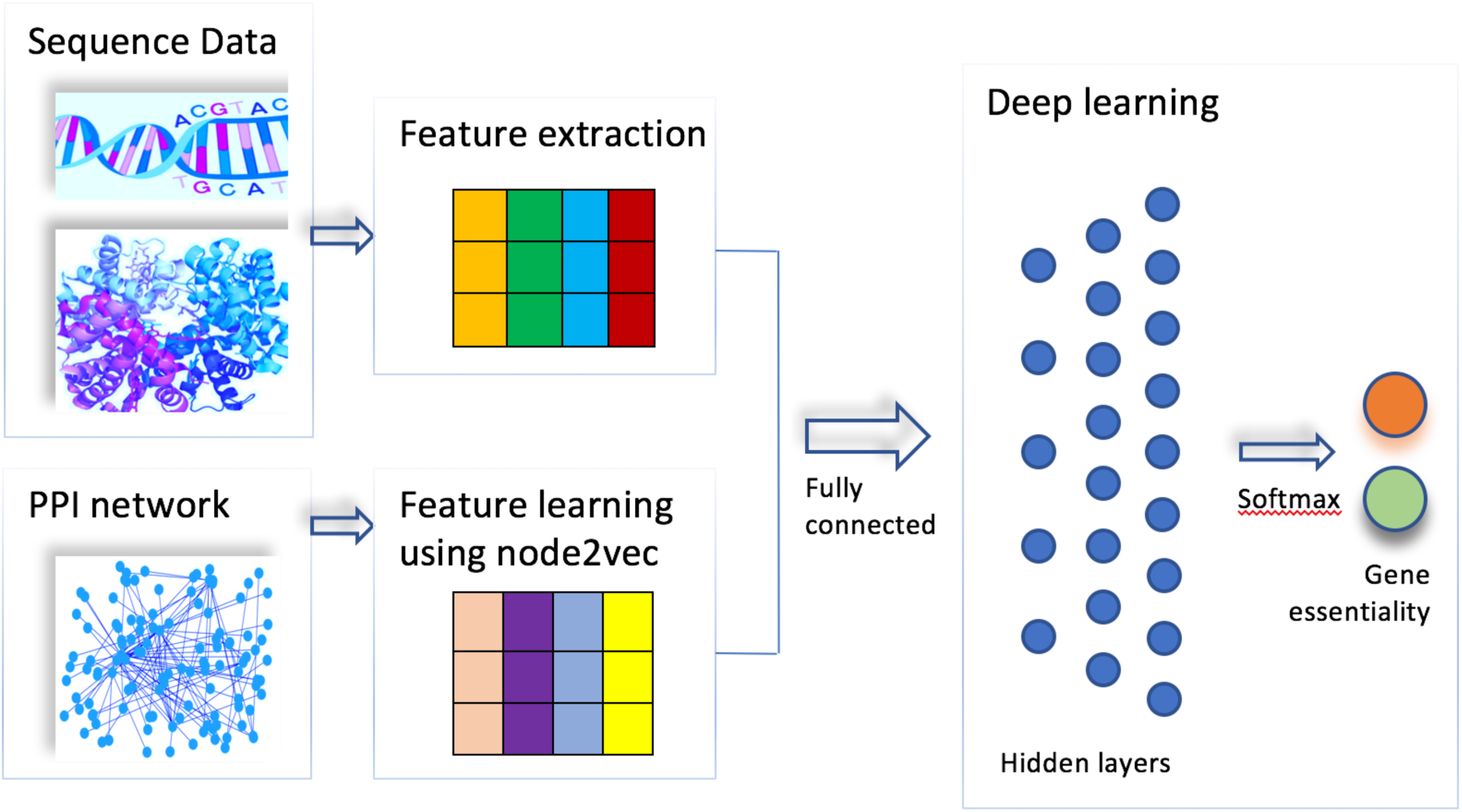
The flowchart of DeepHE.

### 2.1 Features derived from sequence data

We extracted features from gene nucleotide sequences and protein sequences. Several features derived from sequence data have been validated their usefulness in predicting gene essentiality in model organisms [10, 16]. In this paper, we used the following sequence derived features: codon frequency, maximum relative synonymous codon usage (RSCU*max*), codon adaptation index (CAI), gene length, GC content, amino acid frequency, and protein sequence length.

Codon frequency of a gene is computed by sliding a window of three nucleotides along its DNA sequence. The raw counts of 64 codons for each gene were calculated and normalized. Unbalanced synonymous codon usage is prevalent in both prokaryotes and eukaryotes. Codon usage bias in a gene may imply its foreign origin, different functional constraints or a different regional mutation. RSCU is a simple measure of non-uniform usage of synonymous codons in a coding sequence, which is defined as the number of times a particular codon is observed, relative to the number of times that the codon would be observed for a uniform synonymous codon usage. Given a synonymous codon *i* that has an *n*-fold degenerate amino acid, RSCU is computed as (1), where *X*_*i*_ is the number of occurrence of codon *i*, and *n* is 1, 2, 3, 4, or 6 according to the genetic code. In this paper, we used the maximal RSCU of each gene as a feature.

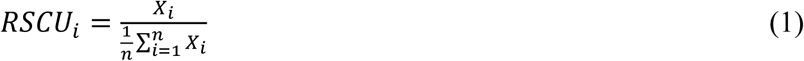

Codon adaptation index (CAI) estimates the bias towards certain codon that are more common in highly expressed genes. The CAI of a gene is defined as (2) where *L* is the number of codons in the gene excluding methionine, tryptophan, and stop codon.

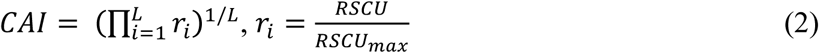

In addition to the 68 features derived from gene nucleotide sequences (64 codon frequency and 1 GC content, gene length, CAI, and RSCUmax, respectively), we also used amino acids frequencies and the protein length, that is, 21 features derived from protein sequences. All features were scaled to have mean *m* = 0 and standard deviation *std* = 1.

### 2.2 Features learned from PPI network

Network embedding methods aim at learning low-dimensional latent representation of nodes in a network, and these representations can be used as features for classification task. Different from some common used topological features, such as node degree centrality (DC), betweenness centrality (BC), and closeness centrality (CC), which usually capture one type of network topological characteristics, the feature representations learned by embedding methods are expected to capture the similarity between nodes in a network.

In this paper, we used a network embedding method, node2vec [15], to automatically learn features for each gene from PPI network. It utilizes a flexible notion of a node’s network neighborhood and a biased random walk procedure to learn richer representations. It aims to learn a mapping of nodes to a low-dimensional space of features that maximizes the likelihood of preserving network neighborhoods of nodes. The biased random walk procedure will generate a corpus which consists of many routes each including multiple nodes. These routes just like the sentences including multiple words in natural language. Then these routes will be fed to word2vec framework using a skip-gram technique to learn low-dimensional features for each node. We got 64 features for each gene from the PPI network.

### 2.3 Deep learning model based on multilayer perceptron

The classification module in our deep learning model, DeepHE, is based on the multilayer perceptron structure. It includes one input layer, three hidden layers, and one output layer. All the hidden layers utilize the rectified linear unit (ReLU) activation function. A ReLU is simply defined as *f*(*x*) = max(0, *x*), which turns negative values to zero and grows linearly for positive values. In DeepHE, the output layer uses sigmoid activation function to perform discrete classification. The loss function in DeepHE is binary cross-entropy.

After each hidden layer, a dropout layer is used to make the network less sensitive to noise in the training data and increase its ability to generalize. The dropout layer randomly assigns zero weights to a fraction of the neurons in the network. Table 1 gives the parameters used in DeepHE.

**Table 1.** Parameters of DeepHE.

|  | #nodes | Activation function | Dropout probability |
| --- | --- | --- | --- |
| Input layer | 153 | - | - |
| Hidden layer 1 | 124 | ReLU | 0.2 |
| Hidden layer 2 | 256 | ReLU | 0.2 |
| Hidden layer 3 | 512 | ReLU | 0.2 |
| Output layer | 2 | sigmoid | - |
| epochs | 100 (early stopping) |  |  |
| optimizer | Adam (learning_rate=0.001) |  |  |

The output *y* of layer *i* depends on the input of layer *i* - 1 as shown in (3), where x is the input, *σ* is the activation function, *b* is the bias, and *W* is the edge weight matrix. During the training phase, the network learns the weights *W* and the bias *b*.

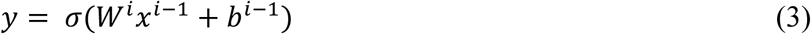

In order to tackle the imbalanced classification problem, we used class weight to train a weighted neural network or cost-sensitive neural network. In the weighted neural network, the backpropagation algorithm will be updated to weigh misclassification errors in proportion to the importance of the class. This will allow the model to pay more attention to examples from the minority class than the majority class in datasets with a severely skewed class distribution.

## 3 Results and discussion

### 3.1 Data collection

DEG database [17] contains 16 human essential gene datasets, among which 13 datasets are from [11-13], and the other three datasets are from [18-20]. We downloaded all the 16 human essential gene datasets for analysis. In total 8,256 human genes are annotated to be essential. Figure 2 shows the distribution of these essential genes across the datasets. According to the assumption that about 10% human genes might be essential genes [12], we selected the genes contained at least in 5 datasets as our essential gene dataset, which has 2,024 genes accounting for ∼10% of human genes. The DNA sequences and protein sequences for essential genes were downloaded from DEG. We downloaded the genome DNA sequences and protein sequences for all annotated genes from Ensembl [21] (release 97, July 2019). Excluding the 8,256 annotated essential genes in DEG, the other annotated protein coding genes formed our nonessential gene dataset, which has 12,697 genes.

**Figure 2.**
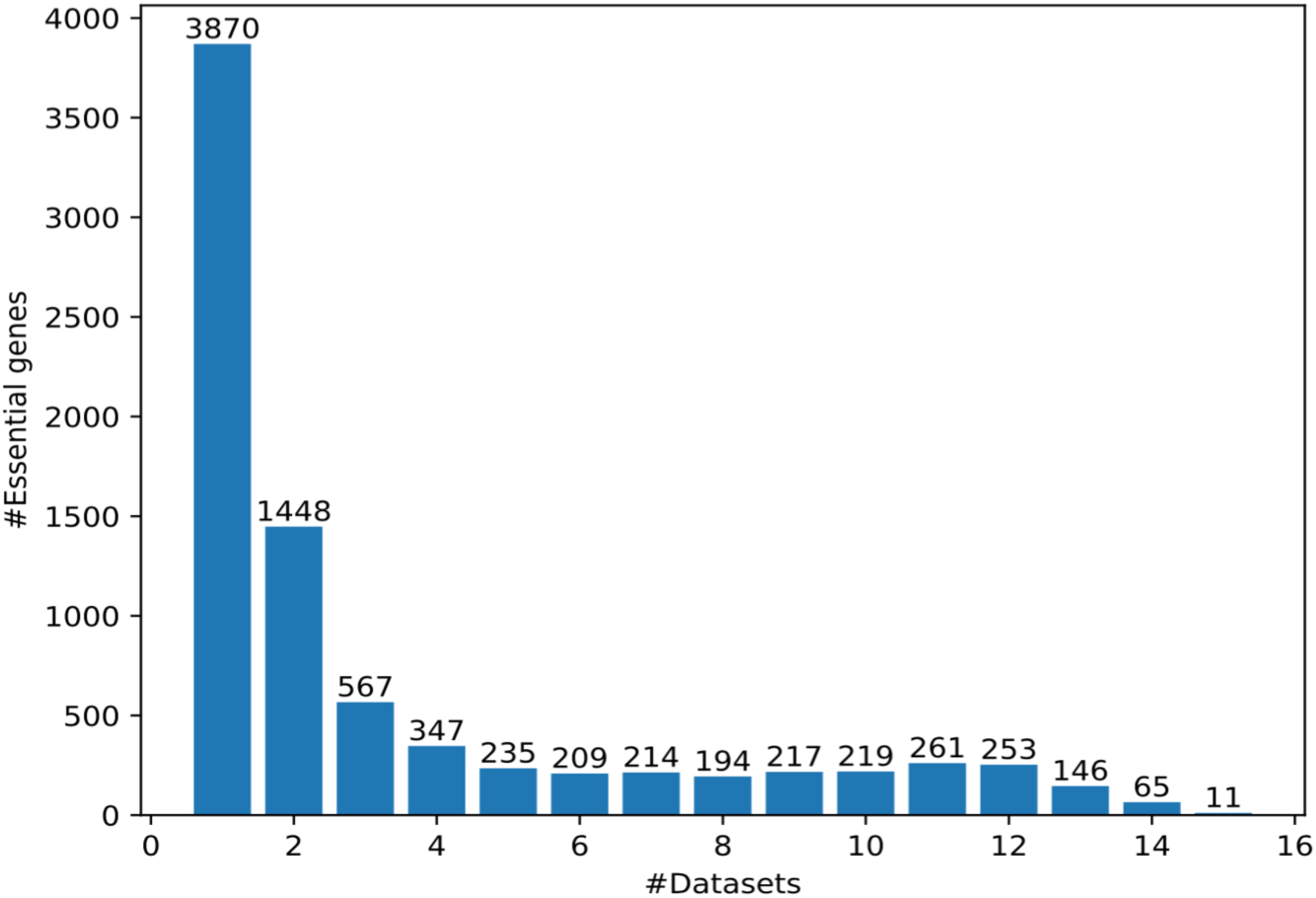
The distribution of essential genes across the 16 datasets.

The protein-protein interaction data was downloaded from BioGRID [22] (release 3.5.181, February 2020). Only physical interactions between human genes were used. After filtering out self-interactions and several small separated subgraphs, we obtained a protein-protein interaction graph with 17,762 nodes and 355,647 edges. This interaction network was used to learn embedding features. We used genes having both sequence features and network embedding features for training and testing the classification model, that is, 2,009 essential genes and 8,430 nonessential genes were used in the following classification performance evaluation.

The number of nonessential genes is more than 4 folds of that of essential genes, which would suffer the class imbalance problem and result in low predictive accuracy issue for the infrequent class. To address this imbalance issue, class weight was used to train a weighted neural network. In each experiment, the 2009 essential genes and 2009 * 4 random selected nonessential genes were used to train, validate and test the model. The class weight was set to 4 for the class of essential genes, and 1 for that of nonessential genes. We also tested the effect of different weights to the performance of our model.

### 3.2 Evaluation metrics

The performance of DeepHE was evaluated using the area under the curve (AUC) of the receiver operating characteristic curve (ROC). ROC plot represents the trade-off between sensitivity and specificity for all possible thresholds. We also used the area under precision-recall curve (AP) to evaluate its performance. Precision-Recall (PR) curves summarize the trade-off between the true positive rate and the positive predictive value for DeepHE using different probability thresholds. ROC curves are appropriate for balanced classification problems in which each class has almost identical number of instances while PR curves are more appropriate for imbalanced datasets. Since human essential gene prediction is an imbalanced classification problem, the area under the PR curve (AP) should be more indicative than AUC-ROC. In addition to AUC and AP scores, we also gave the following performance measures: sensitivity (Sn), specificity (Sp), positive predictive value (PPV), and accuracy (Ac), which are defined in (4) - (7), where *TP, TN, FP*, and *FN* are the number of true positives, true negatives, false positives, and false negatives, respectively.

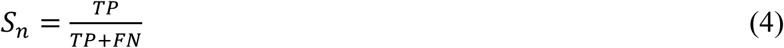

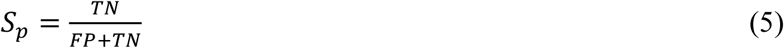

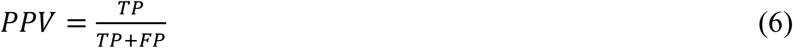

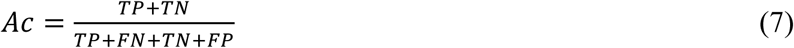

### 3.3 Performance evaluation

#### 3.3.1 The effect of number of hidden layers and dropout probability

There are several hyper-parameters in DeepHE, which would affect its performance. In the following experiments, we chose Adam as the optimizer because of its superior performance. Its initial learning rate is 0.001. The training was run for 100 epochs with early stopping criteria. The batch size is 32. For each run, the 2009 * 4 nonessential genes were randomly selected from the 8430 nonessential genes. We used 80% data for training, 10% data for validation, and the other 10% data for testing. We kept the same ratio between the number of essential genes and that of nonessential genes in training, validating, and testing data. Each experiment was executed 10 times to get the average performance.

Table 2 gives the performance of DeepHE with different number of hidden layers and different dropout probability (DP). From table 2 we can see that the overall performance of DeepHE is very robust to these two parameters. For example, its best, average, and worst AUC scores are 94.15%, 93.23%, and 92.47% respectively. It achieves the best overall performance with AUC = 94.15% when using HL3 with DP = 0.2. Its AP scores are also very stable with the best, average, and worst values of 90.64%, 89.4%, and 88.69% respectively. Same with AUC, it achieves the best AP score of 90.64% when using HL3 with DP = 0.2. In addition to the best AUC and AP scores, it also achieves the best scores for specificity (94.5%), PPV (77.74%), and accuracy (90.88%) when using HL3 with DP = 0.2. The best sensitivity score is 87.16% when using HL5 with DP = 0.5. From table 2 we can also see that with the increase of drop probability, its sensitivity score increases but its PPV score decreases in most cases.

**Table 2.** The performance of DeepHE with different parameters.

| Hidden layers | Dropout probability | AUC | Sn | Sp | PPV | Ac | AP |
| --- | --- | --- | --- | --- | --- | --- | --- |
| HL3<br>[128, 256, 512] | 0.1 | 0.9325 | 0.7537 | 0.9295 | 0.7303 | 0.8943 | 0.8882 |
|  | 0.2 | **0.9415** | 0.7638 | **0.945** | **0.7774** | **0.9088** | **0.9064** |
|  | 0.3 | 0.9352 | 0.7716 | 0.928 | 0.7302 | 0.8967 | 0.8967 |
|  | 0.4 | 0.9356 | 0.7881 | 0.9188 | 0.7091 | 0.8926 | 0.8972 |
|  | 0.5 | 0.9321 | 0.808 | 0.8937 | 0.6571 | 0.8765 | 0.8904 |
| HL4<br>[128, 256, 512, 1024] | 0.1 | 0.931 | 0.7572 | 0.9295 | 0.7305 | 0.895 | 0.8924 |
|  | 0.2 | 0.9338 | 0.7816 | 0.9235 | 0.7215 | 0.8951 | 0.8976 |
|  | 0.3 | 0.9282 | 0.8025 | 0.9075 | 0.6877 | 0.8865 | 0.8942 |
|  | 0.4 | 0.9372 | 0.8488 | 0.8711 | 0.624 | 0.8667 | 0.8967 |
|  | 0.5 | 0.9369 | 0.8687 | 0.8553 | 0.601 | 0.858 | 0.9018 |
| HL5<br>[128, 256, 512, 1024, 1024] | 0.1 | 0.9247 | 0.7672 | 0.921 | 0.711 | 0.8902 | 0.8888 |
|  | 0.2 | 0.931 | 0.7811 | 0.9211 | 0.7129 | 0.8931 | 0.893 |
|  | 0.3 | 0.9282 | 0.8199 | 0.8893 | 0.6515 | 0.8754 | 0.8891 |
|  | 0.4 | 0.9283 | 0.8498 | 0.8654 | 0.6128 | 0.8623 | 0.8909 |
|  | 0.5 | 0.9296 | **0.8716** | 0.827 | 0.5583 | 0.8359 | 0.8869 |

In a very skewed classification problem, the accuracy and AUC measures can get large values even when almost all the instances in the minority class are classified into the majority class. That’s not what we expected. In most cases of imbalanced classification problems, we are far more concerned with the classifier’s performance on the minority class. Since the essential gene prediction problem is often a very skewed classification problem in which the number of essential genes is much less than that of nonessential genes. Our concerns would be how many essential genes can be predicted and how many genes are truly essential among those predicted as essential genes, that is, sensitivity and PPV as well as the comprehensive measure AP are more important. Based on this point, we think that DeepHE with 3 hidden layers and DP = 0.2 is the best one which will be used in the following experiments.

Figure 3 gives the ROC curves of DeepHE in 10 repetitions when using HL3 and DP = 0.2. ROC curves summarize the trade-off between the true positive rate and false positive rate of DeepHE using different probability thresholds. From figure 3 we can see that DeepHE reached its best performance at iterations 2, 4, 8, and 10 with AUC = 0.95. In addition, the performance of DeepHE is quite stable since the difference is only about 0.02 between its best and worst AUC scores. Guo et al. also used machine learning (SVM) to predict human essential genes based on sequence data [8]. Their reported best performance is AUC = 0.88. Compared with [8], DeepHE outperformed their method.

**Figure 3.**
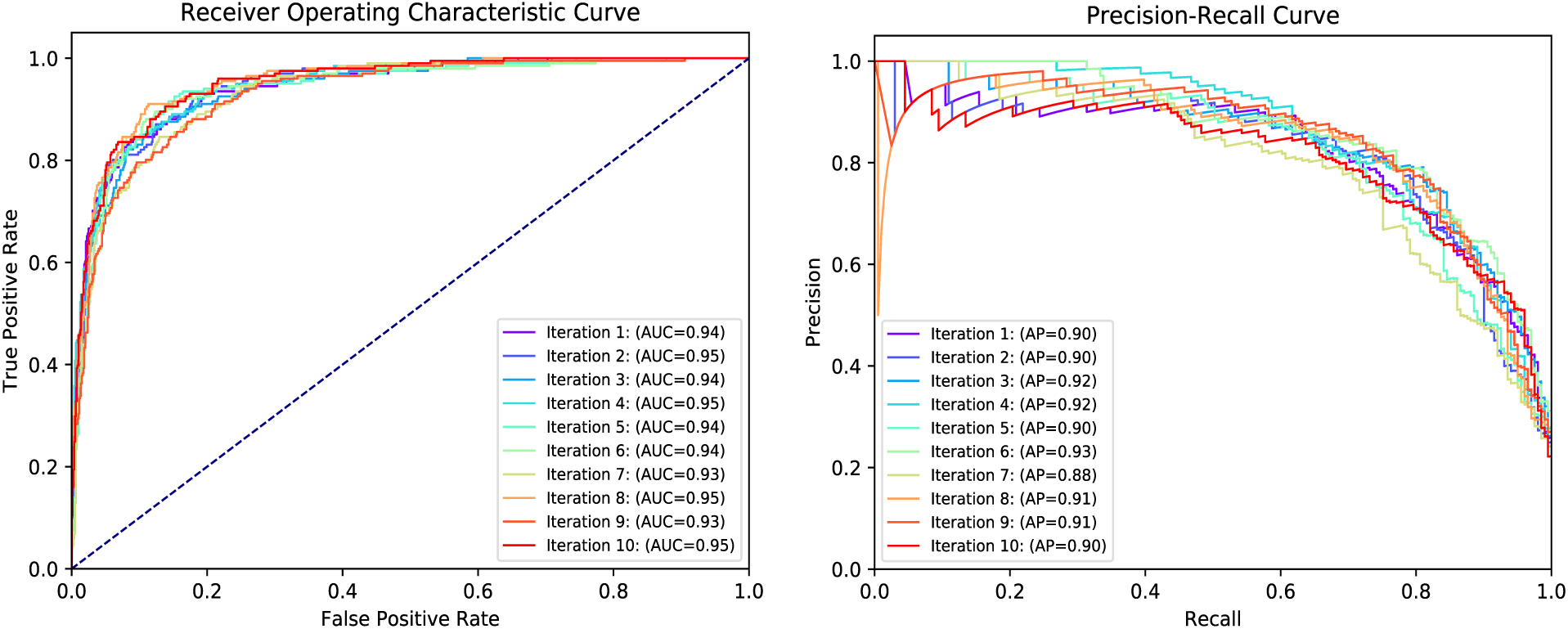
The ROC and PR curves of DeepHE with HL3 and DP = 0.2.

Figure 3 also shows PR curves for 10 iterations of DeepHE with HL3 and DP = 0.3. Similar with the AUC scores, its AP scores are also very stable since there’s only a very small difference between its best and worst AP scores (about 0.04). It achieves the best performance in iteration 6 with AP = 93%. The worst AP score is still above 88% which indicates that DeepHE is very effective for predicting human essential genes.

#### 3.3.2 The effect of class weight

In order to cope with the imbalanced data distributions between two classes, DeepHE used class weight to give larger penalty when misclassifying an instance in the minority class, that is, the class of essential genes. In the following, we will test if different weight values would affect the performance of DeepHE. Note that in each experiment, the ratio between the number of essential genes and that of nonessential genes is 1:4. The class weight for nonessential genes is always 1. We will vary the class weight for essential genes from 1 to 10 to see its effect on the performance. DeepHE with 3 hidden layers and DP = 0.2 is used for the following experiments.

Table 3 gives the performance of DeepHE with different class weights for the class of essential genes. From table 3 we can see that DeepHE achieves best AUC (94.15%), PPV (77.74%), Accuracy (90.88%), and AP score (90.64%) when class weight = 4.0. It gets the best sensitivity score (79.85%) when class weight = 9.0. In general, the sensitivity score increases with the increase of the class weight, but PPV score decreases with the increase of class weight. This accords with our intuition. With larger class weight, misclassifying an essential gene will get larger penalty than misclassifying a nonessential gene. In this situation, more essential genes will be put into the right class, at the same time, more nonessential genes would also be put into the class of essential genes, which will result in higher sensitivity score and lower PPV score. When class weight = 4.0, it mimics the situation that the number of essential genes equal to the number of nonessential genes, thus it achieves a balanced point for sensitivity and PPV score. One can set the class weight according to his preference on whether higher specificity or higher PPV or just the balance between them.

**Table 3.** The performance of DeepHE with different class weights for essential genes.

| Class weight | AUC | Sn | Sp | PPV | Ac | AP |
| --- | --- | --- | --- | --- | --- | --- |
| 1.0 | 0.9307 | 0.7189 | **0.9475** | 0.7744 | 0.9018 | 0.8949 |
| 2.0 | 0.9321 | 0.7328 | 0.9377 | 0.7462 | 0.8967 | 0.8934 |
| 3.0 | 0.9368 | 0.7438 | 0.9388 | 0.7524 | 0.8998 | 0.8969 |
| 4.0 | **0.9415** | 0.7638 | 0.945 | **0.7774** | **0.9088** | **0.9064** |
| 5.0 | 0.9369 | 0.7796 | 0.9261 | 0.7288 | 0.8968 | 0.8982 |
| 6.0 | 0.9349 | 0.7701 | 0.9284 | 0.7301 | 0.8967 | 0.8963 |
| 7.0 | 0.9322 | 0.7627 | 0.9275 | 0.7253 | 0.8945 | 0.8919 |
| 8.0 | 0.9336 | 0.7841 | 0.9215 | 0.716 | 0.894 | 0.8957 |
| 9.0 | 0.9381 | **0.7985** | 0.9209 | 0.7173 | 0.8964 | 0.9031 |
| 10.0 | 0.9316 | 0.7816 | 0.9095 | 0.6851 | 0.8839 | 0.8922 |

Table 3 also tells us that the AUC, specificity, AP, and Accuracy of DeepHE are very robust to the class weight. For example, the best, average, and worst AUC scores are 94.15%, 93.48%, and 93.07% respectively; the best, average, and worst specificity scores are 94.75%, 93.03%, and 90.95% respectively; the best, average, and worst AP scores are 90.64%, 89.69%, and 89.19% respectively; the best, average, and worst Accuracy scores are 90.88%, 89.69%, and 88.39% respectively. When varying the class weight, sensitivity score and PPV change in opposite directions which makes the overall performance of DeepHE only slightly affected by the change of class weight.

#### 3.3.3 The contribution of different features

DeepHE utilizes two types of features, sequence features (S) and network embedding features (N). In the following we will test how each type of features affect the performance of DeepHE. In the following experiments, DeepHE works with same configurations (3 hidden layers, DP = 0.2, class weight = 4.0. Other configurations are same as before) except the input features.

Table 4 gives the performance of DeepHE using different type of features. It tells us that DeepHE with the integration of sequence features and network embedding features works best which confirms the contribution of the two types of features. DeepHE with only sequence features works worst which has very low PPV score (53.28%). DeepHE with network embedding features works in between, whose AP score achieves acceptable level (86.53%). DeepHE achieves the best performance for all the six measures by integrating these two types of features.

**Table 4.**
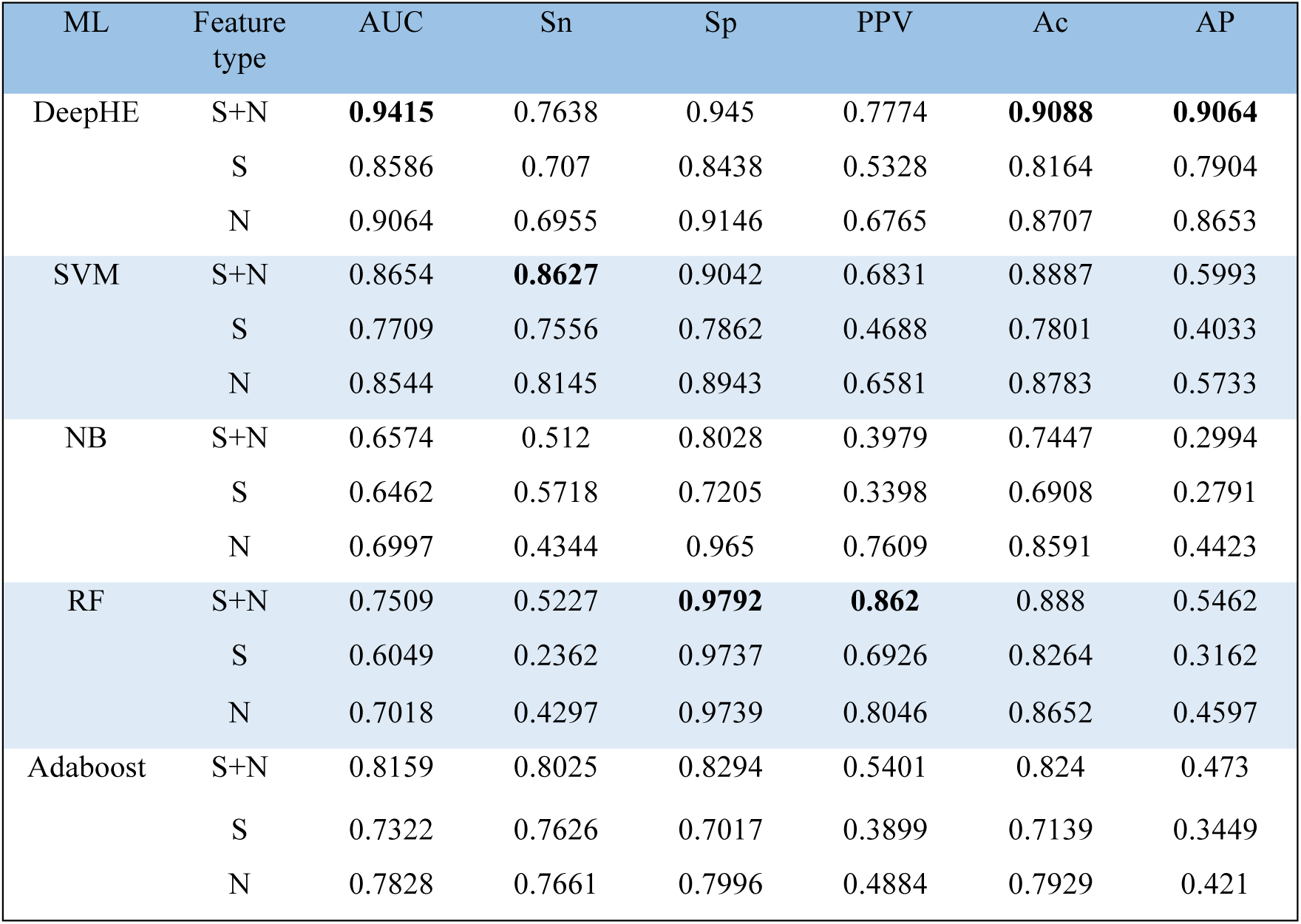
The performance of DeepHE, SVM, NB, RF, and Adaboost with different type of features.

#### 3.3.4 Comparison with traditional machine learning models

Several machine learning methods have been used to predict essential genes [7]. In order to demonstrate the superior of our proposed prediction method DeepHE, we also compared it with several widely used traditional machine learning models, such as Support Vector Machines (SVM), Naïve Bayes (NB), Random Forest (RF), and Adaboost. All the compared machine learning algorithms are implemented by scikit-learn python library with default parameters, unless otherwise specified. For each model, we either set class_weight parameter to 4.0 or set sample_weight parameter to 4.0 for each essential gene and 1.0 for each nonessential gene, therefore the two types of weights are essentially same. The sample_weight is only used when class_weight is not available. All models were tested 10 times and the average performance for each measure was reported.

Table 4 gives the results. From table 4 we can see that DeepHE (N+S) significantly outperforms the other machine learning models regarding to three comprehensive measures, AUC, AP, and Accuracy. For instance, the AUC score of DeepHE (N+S) is 8.79% higher than that of SVM (N+S), 43.22% higher than that of NB (N+S), 25.38% higher than that of RF (N+S), and 15.39% higher than that of Adaboost (N+S). The AP score of DeepHE (N+S) increases by 51.24%, 220.77%, 65.95%, and 91.63% compared with that of SVM (N+S), NB (N+S), RF (N+S), and Adaboost (N+S) respectively. By integrating sequence features and network embedding features, the overall performance of four models (DeepHE, SVM, RF, Adaboost) gets improved. NB works slightly better with only network embedding features. Considering the fact that essential gene prediction is an imbalanced problem, AP is more important than other measures. From table 4 we can see that the four compared machine learning models all have very low AP scores (from 27.91% to 59.93%) which tells us that they are not a good choice for such task, and further confirms the superior of our proposed deep learning model, DeepHE.

## 4 Conclusion

We proposed a new essential gene prediction framework based on deep learning, DeepHE. It aims to explore whether deep learning can achieve notable improvements for predicting gene essentiality, an imbalanced classification problem. DeepHE integrates two types of features, sequence features extracted from DNA sequence and protein sequence and features learned from PPI network, as its input. Then a multilayer perceptron was used to train a cost-sensitive classifier by setting class weight. Although several machine learning based essential gene prediction methods have been proposed, most of them based on the features extracted according to human domain knowledge. In this paper, we used a deep learning model, node2vec, to automatically learn network features for each gene from the PPI network. The learned embedding features greatly improved the performance of DeepHE compared with it only using sequence features. The performance of DeepHE was evaluated on human datasets, which achieved very good performance for three comprehensive measures AUC (94.15%), AP (90.64%), and Accuracy (90.88%). We also compared it with four widely used machine learning models, SVM, Naïve Bayes, Random Forest, and Adaboost. DeepHE significantly outperforms all the four machine learning models, which further demonstrates that DeepHE is an effective deep learning framework for human essential gene prediction.

In the future, we will explore other biological data to further improve the performance of DeepHE. Especially we are interested in how to use deep learning to automatically learn features from biological data rather than manually extracting features heavily based on domain knowledge. In addition, we are also interested in exploring more useful techniques to cope with the imbalanced classification problem as well as sparsely labeled classification problem [23,24]. Exploring deep learning to predict human essential genes across human cancer cell lines would be also interesting.

## Authors’ contributions

XZ conceived and designed the experiments. XZ and WJX performed the experiments. XZ and WXX drafted and revised the manuscript. All authors approved the final manuscript.

## Data Availability

All data used in this study are third party and freely accessible from public databases. Protein-protein interaction data are available from BioGRID database at http://thebiogrid.org/download.php. Essential genes data and the corresponding sequence data from DEG database are available at http://tubic.tju.edu.cn/deg/. DNA sequence and protein sequence data are available at https://useast.ensembl.org/Homo_sapiens/Info/Annotation.

## References

1. Zhang X, Xu J, Xiao W-x. A New Method for the Discovery of Essential Proteins. PLoS ONE. 2013; 8 (3): e58763. https://doi.org/10.1371/journal.pone.0058763.

2. Zhang X, Xiao W, Acencio ML, Lemke N, Wang X. An ensemble framework for identifying essential proteins. BMC Bioinformatics. 2016; 17:322. https://doi.org/10.1186/s12859-016-1166-7.

3. Zhang X, Xiao W, Hu X. Predicting essential proteins by integrating orthology, gene expressions, and PPI networks. PLoS ONE, 2018, 13(4): e0195410. https://doi.org/10.1371/journal.pone.0195410.

4. Li G, Li M, Wang J, Wu J, Wu F, Pan Y. Predicting essential proteins based on subcellular localization, orthology and PPI networks. BMC Bioinformatics. 2016; 17(Suppl 8):279. https://doi.org/10.1186/s12859-016-1115-5.

5. Peng W, Wang J, Cheng Y, Lu Y, Wu F, Pan Y. UDoNC: an algorithm for identifying essential proteins based on protein domains and protein-protein interaction networks. IEEE/ACM Transactions on Computational Biology and Bioinformatics, 12(2): 276–288, 2015.

6. Li X, Li W, Zeng M, Zheng R, Li M. Network-based methods for predicting essential genes or proteins: a survey. Briefings in Bioinformatics, bbz017, 2019. https:doi.org/10.1093/bib/bbz017.

7. Zhang X, Acencio ML and Lemke N. Predicting Essential Genes and Proteins Based on Machine Learning and Network Topological Features: A Comprehensive Review. Front. Physiol. 2016; 7:75. https://doi.org/10.3389/fphys.2016.00075 PMID: 27014079.

8. Guo F, Dong C, Hua H, Liu S, Luo H, Zhang H, Jin Y, Zhang K. Accurate prediction of human essential genes using only nucleotide composition and association information. Bioinformatics, 33(12), 2017, 1758–1764. doi: 10.1093/bioinformatics/btx055.

9. Zeng M, Li M, Fei Z, Wu F, Li Y, Pan Y, Wang J. A deep learning framework for identifying essential proteins by integrating multiple types of biological information. IEEE/ACM Transactions on Computational Biology and Bioinformatics. 2019 Feb 5. doi: 10.1109/TCBB.2019.2897679.

10. Hasan MA, Lonardi S. DEEPLYESSENTIAL: A deep neural network for predicting essential genes in microbes. BioRxiv, 2019. http://dx.doi.org/10.1101/607085.

11. Blomen VA, Májek P, Jae LT, Bigenzahn JW, Nieuwenhuis J, et al. Gene essentiality and synthetic lethality in haploid human cells. Science, 2015, 350(6264):1092–6. doi: 10.1126/science.aac7557.

12. Wang T, Birsoy K, Hughes NW, Krupczak KM, Post Y, et al. Identification and characterization of essential genes in the human genome. Science, 2015, 350(6264): 1096–101. doi: 10.1126/science.aac7041.

13. Hart T, Chandrashekhar M, Aregger M, Steinhart Z, Brown KR, et al. High-resolution CRISPR screens reveal fitness genes and genotype-specific cancer liabilities. Cell, 2015, 163(6): 1515–26. doi:10.1016/j.cell.2015.11.015.

14. Fraser A. Essential human genes. Cell Systems, 2015, 1(6): 381–382. doi: 10.1016/j.cels.2015.12.007.

15. Grover A, Leskovec J (2016). node2vec: Scalable Feature learning from networks. KDD’16: Proceedings of the 22^nd^ ACM SIGKDD International Conference on Knowledge Discovery and Data Mining. August 2016, pp 855–864. https://doi.org/10.1145/2939672.2939754

16. Liu X, Wang BJ, Xu L, Tang HL, Xu GQ (2017) Selection of key sequence-based features for prediction of essential genes in 31 diverse bacterial species. PLoS ONE 12(3): e0174638. https://doi.org/10.1371/journal.pone.0174638.

17. Luo H, Lin Y, Gao F, Zhang CT, and Zhang R. DEG 10, an update of the database of essential genes that includes both protein-coding genes and noncoding genomic elements. Nucleic Acids Res., 42 (Database issue): D574–80, Jan. 2014.

18. Liao BY, Zhang J (2008). Null mutations in human and mouse orthologs frequently result in different phenotypes. Proc Natl Acad Sci U S A, 105:6987-92.

19. Georgi, Benjamin, Benjamin F. Voight, and Maja Bucan. From Mouse to Human: Evolutionary Genomics Analysis of Human Orthologs of Essential Genes. PLoS genetics 9.5 (2013): e1003484

20. Lek, Monkol, et al. Analysis of protein-coding genetic variation in 60,706 humans. bioRxiv (2015): 030338.

21. Ruffier M, Kähäri A, Komorowska M, Keenan S, Laird M, et al. Ensembl core software resources: storage and programmatic access for DNA sequence and genome annotation. Database, 2017, doi: 10.1093/database/bax020.

22. Stark C, Breitkreutz BJ, Reguly T, Boucher L, Breitkreutz A, Tyers M. Biogrid: A General Repository for Interaction Datasets. Nucleic Acids Res. Jan 1, 2006; 34: D535–9.

23. Zhang X, Xiao W. Clustering based two-stage text classification requiring minimal training data. Computer Science and Information Systems. 2012; 9(4):1627–1643. doi: 10.2298/CSIS120130044Z

24. Zhang X, Xiao W. Active semi-supervised framework with data editing. Computer Science and Information Systems. 2012; 9(4): 1513–1532. doi: 10.2298/CSIS120202045Z

